# Morphometric Identification of Stem Borers *Diatraea saccharalis* and *Diatraea busckella* (Lepidoptera: Crambidae) in Sugarcane Crops (*Saccharum officinarum*) in Caldas Department, Colombia

**DOI:** 10.1101/385955

**Authors:** Julián Andrés Valencia Arbeláez, Alberto Soto Giraldo, Gabriel Jaime Castaño Villa, Luis Fernando Vallejo Espinosa, Melba Ruth Salazar Guitierrez^4^

## Abstract

The sugarcane (*Saccharum* sp.), of great importance for being one of the most traditional rural agroindustries in Latin America and the Caribbean, as part of the agricultural systems, is vulnerable to increases or reductions in the incidence of pests associated with extreme events of climate change, such as prolonged droughts, hurricanes, heavy and out of season rains, among others, contributing to the increase losses in agricultural production, which forces farmers to make excessive expenditures on pesticides that generally fail to solve the issue. (Vázquez, 2011). The main pest belongs to the *Diatraea* complex (Vargas et al., 2013; Gallego *et al.,* 1996), a larval stage perforator habit. Different field evaluations have revealed the presence of a species that had not been reported in sugarcane crops, *Diatraea busckella*, and to corroborate the finding, a method of identification was needed whose advantage was to be quick and also low cost, in this sense, geometric morphometry is a mathematical tool with biological basis (Bookstein, 1991), which allows to decompose the variation resulting from the physiology of individuals of the most stable individuals of the population, product of the genetic component. CLIC (Collecting Landmarks for Identification and Characterization) was used for identification, with reference to the previous right wing (De La Riva *et al*., 2001; Belen *et al*., 2004; Schachter-Broide *et al*., 2004; Dvorak *et al*., 2006; Soto Vivas *et al.,* 2007). Wing morphometry was performed using generalized Procrustes analysis (Rohlf and Marcus, 1993). The analysis clearly differentiated between *D. busckella* and *D. saccharalis*, eliminating the environmental factors that could generate some level of error, being considered a support tool that validates the molecular biology processes for the identification of organisms.

## Introduction

Recent studies have highlighted the vulnerability of agriculture to climate variables and the effects on harvests of pests and diseases (Lau *et al*., 2011; Carvajal, 2010; Contreras and Galindo, 2009). Within this context, in agricultural systems, it has been possible to verify increases or reductions in the incidence of pests associated with extreme events of climate change, such as prolonged droughts, hurricanes, heavy and out of season rains, among others. Of course, these are often not measurable, because the disasters caused by such events in crops do not allow us to appreciate the changes in the manifestations of pests. However, these contribute to increased losses, forcing farmers to spend too much on pesticides that usually fail to solve the problem. (Vázquez, 2011).

A crop that is not exempt from these changes of climatic variability is the sugarcane (*Saccharum* sp.), of great importance for being one of the most traditional rural agroindustries in Latin America and the Caribbean and whose world production is around 13 million tons per year (FAO, 2007). In order of importance, the first country to produce sugarcane is India, followed by Colombia, where it is the second largest source of rural economic production after coffee (CENICAÑA, 2007; DANE, 2007; FAO, 2007). As with all agricultural holding, sugarcane crop is no stranger to pest attacks, with the main impact being the borers of the *Diatraea* complex (Vargas *et al*., 2013; Gallego *et al*., 2007), 1996), and its permanent presence in the larval stages and its permanent presence either in the mouths or in new crops, causes from damage to the bud in the initial stages of the crop, to perforations in the nodes or internodes causing the plant to tip over and loss in quality due to loss of sucrose (Parody, 2011; Gallego *et al*., 1996).

However, numerous studies in different aspects of biology are being carried out, which seek to understand the origin, nature and causes of change in variation patterns, with the aim of finding answers to the integrated management of the insect and control methods, using quantitative analyses of phenotypic variation and its relationship with the environment, genetics and random factors (or of unknown origin), necessary to identify organisms (Jaramillo, 2011).

These insects, like all living organisms, visibly highlight their morphological variability. A number of biological and non-biological factors underlie such variability by interacting in a complex way to model individuals, populations and species (Calle, 2008).

Although the size and colour of larvae and adults are characteristics that, in a very approximate way, allow us to distinguish species of the *Diatraea* spp complex with a relatively low margin of error, more precise and state-specific characteristics are required to identify the species with a high degree of precision (Gómez *et al*., 2014), which is why we mainly use external morphological and genitalia characteristics (Cadena, 2008; Linares and Bastidas, 1996).

These biological factors are the product of micro- and macro-evolutionary processes interacting with the environment. Thus, the size and morphological conformation results from the interaction between maternal effects and quality of life during development with the genotypic combinations inherited from the parents, within a genetic framework given by evolutionary history. The populations and species, then, acquire their own morphological features that have a differential impact on their biology and ecology (Jaramillo, 2011). Evidence of these factors was found in late 2012, near the town of Cartago, Valle del Cauca, Colombia, where *Diatraea* larvae were collected that apparently did not match the characteristics of *D. saccharalis* and *D. indigenella*. Morphological observation of the different stages of life (larva, pupa and adult) allowed the identification of the larvae belonging to the species *Diatraea busckella*, with the initial records of this species attacking sugarcane crops. (Vargas *et al*., 2013).

To corroborate that, a identification method was needed that was both quick and cost-effective. In this sense, geometric morphometry is a mathematical tool with biological grounds (Bookstein, 1991), which allows us to decompose the variation resulting from the physiology of the individuals of the most stable individuals in the population, a product of the genetic component. It allows the identification of individuals from the conformation of specific structures, in closely related species (Dujardin, 2000), based on methods of description and statistical analysis of the form variation in particular structures within and between individuals (Rohlf and Marcus, 1993). This technique has been used in various types of plant and animal organisms (vertebrates and invertebrates), and particularly in insects of agricultural and medical importance, such as *Nymphalidae*, *Tortricidae*, *Noctuidae*, *Triatominae*, and *Culicidae*. It has been used in studies of population structure (De La Riva *et al*., 2001; Belen *et al*., 2004; Schachter-Broide *et al*., 2004; Dvorak *et al.,* 2006; Soto Vivas *et al.,* 2007), species identification (Matías *et al*., 2001; Villegas *et al*., 2002; Calle *et al*., 2008), to detect differences between field parents and laboratory descendants (Jaramillo and Wolff, 2002), as well as in phylogenetic studies (Soto Vivas *et al*., 2011).

Quantifying and analyzing the morphological variation resulting from such interactions is the main objective of morphometry. (Jaramillo, 2011), in addition, there is little research work on aspects of insect identification, particularly related to the use of morphometry.

## Materials and methods

The collections were carried out in 4 sugarcane municipalities in the department of Caldas, Colombia (Table 1) every two weeks in 120 randomly selected points (canes), making a 40 minutes zigzag trip per plot within the cane crops, following the orientation according to the linear distribution of the “chorrillo” (Pérez *et al.,* 2011), or at the time of cutting in the sugarcane mills where samples of eggs, larvae, pupae and adults were taken, from the time when the first true leaves appear in the crop of sugarcane until the time of harvest. The larval collections were manual in each of the sampled sites, in order to obtain the infestation index, recorded as the location of the collection points, so that a representative sampling of the area can be obtained.

**Table 1.**
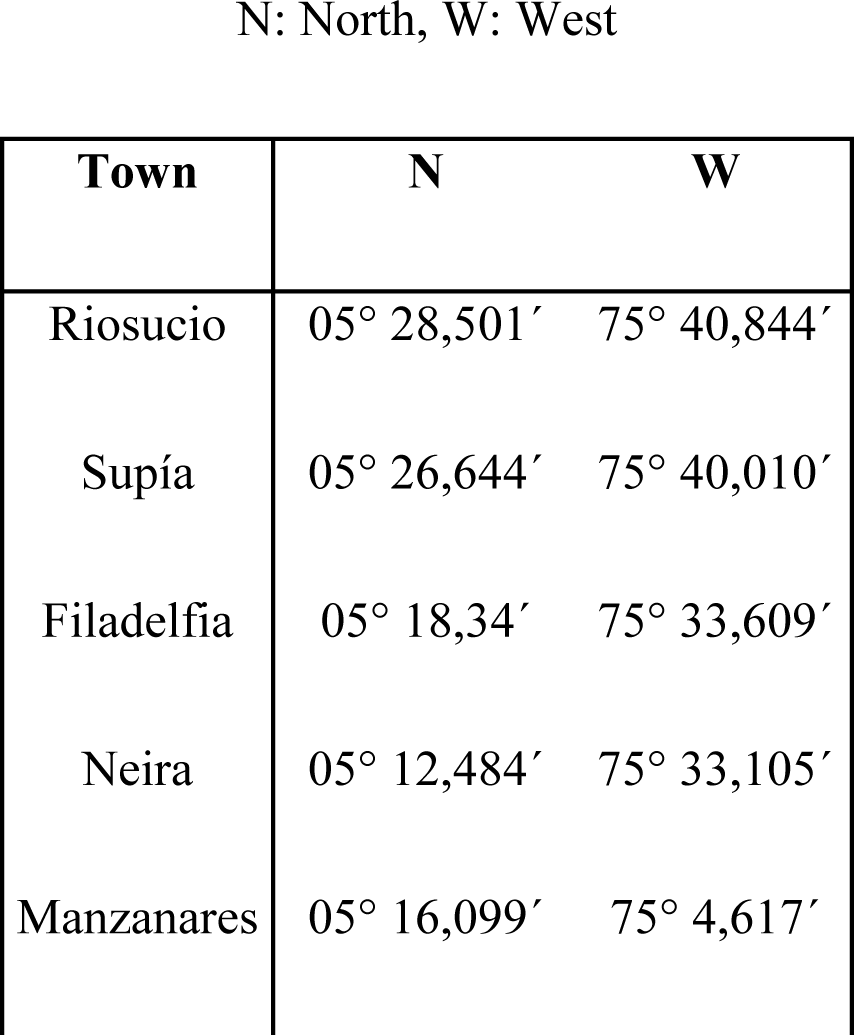
*Geographical location of collection points for* D. busckella *and* D. saccharalis.

N: North, W: West
| Town | N | W |
| --- | --- | --- |
| Riosucio | 05° 28,501' | 75° 40,844' |
| Supía | 05° 26,644' | 75° 40,010' |
| Filadelfia | 05° 18,34' | 75° 33,609' |
| Neira | 05° 12,484' | 75° 33,105' |
| Manzanares | 05° 16,099' | 75° 4,617' |

Each individual collected in the larval stage, was taken to adult, under controlled conditions of temperature (20; 24.7 and 30°C) and relative humidity (35%), using a diet of sweet corn not exceeding 30 days, in the Center for Research and Breeding of Natural Enemies, located in the botanical garden of the University of Caldas at 5° 3' 23''' latitude north and 75° 29' 39''' west longitude, at an altitude of 2160 meters above sea level and with an average temperature of 14°C (Manizales, Caldas).

In order to identify which species are present within each evaluated site, the external morphology of the right rear wing was characterized in an adult state from the junction of the basal portion articulated to the thorax. (Jaramillo, 2011; Cadena, 2008; Pastrana, 1991).

To assemble the wings, the scales were first removed by washing with 10% NaClO for 5 minutes, then rinsed with distilled water for another 5 minutes. The wings were mounted on slide film and for their preservation clear varnish was used and the images were taken from a Nikon SMZ1270 stereoscope at a 1. 5x zoom.

On the wing images, twenty-four (24) points were selected between the limits of the basal and subcostal veins and the intersections of the Cu 1 and 2 veins with the distal edge of the wing (Fig 1 and Fig 2). These points allowed the geometric configuration of each wing to be captured (Calle *et al*., 2008).

**Fig 1.**
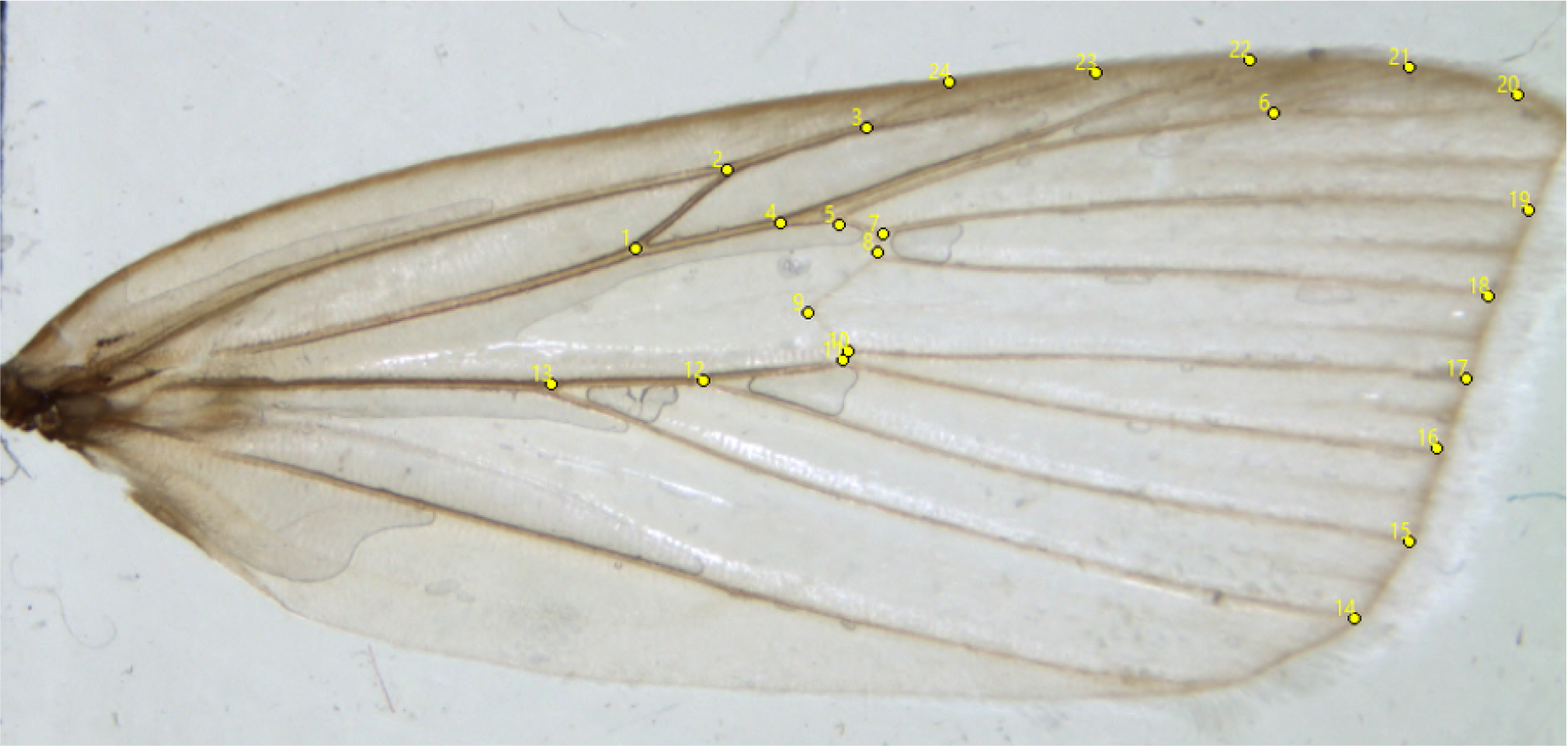
Right rear wing of *D. busckella*. Landmarks for identification of morphometric points.

**Fig 2.**
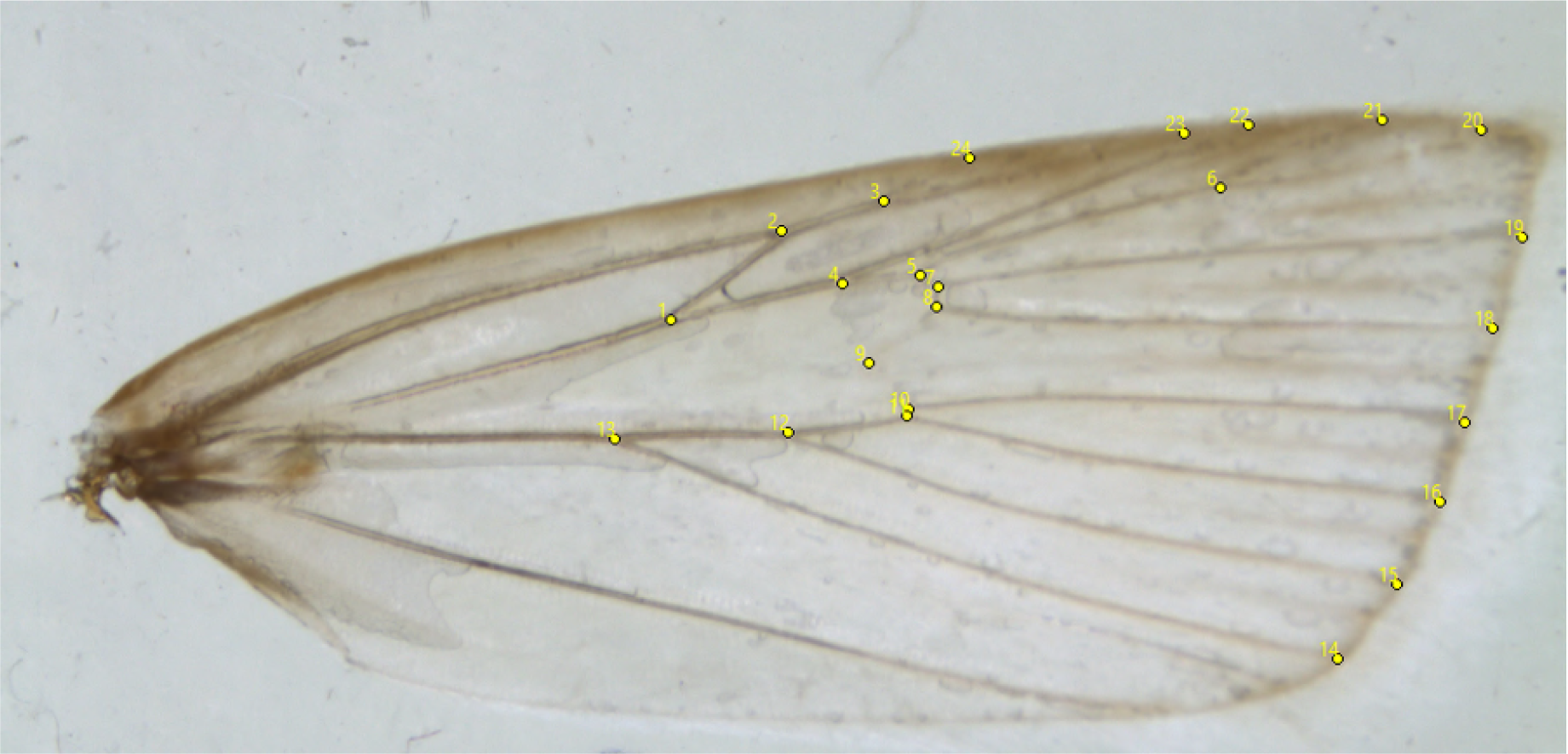
Right rear wing of *D. saccharalis*. Landmarks for identification of morphometric points.

CLIC (Collecting Landmarks for Identification and Characterization) open access software (mome-clic. com) was used for identification, taking as a reference the previous right wing, based on similar research for different arthropods (De La Riva *et al*., 2001; Belen *et al*., 2004; Schachter-Broide *et al*., 2004; Dvorak *et al*., 2006; Soto Vivas *et al*., 2007).

Wing morphometry was performed using generalized Procrustes analysis (Rohlf. And Marcus, 1993; The anatomical reference points were chosen taking into account their easy identification and reproducibility, hence, that they have homology of position among the individuals in the stock (Bookstein, 1991). 24 anatomical reference points were selected. According to their categories (Bookstein, 1991), on the wings all anatomical reference points are type 1.

The geometric configuration of the wings was obtained by capturing the points at the intersections of the radial (R2 and R3), medial (M2 and M3) and cubital (Cu1 and Cu2) veins, with the distal borders: R5, M1, M2, M3, Cu1 and Cu2. (Figures 3 and 4). Based on the Bookstein classification, 1991, all APRs (Anatomical Position of the Reference points) were type I, representing vein-to-vein intersections and vein completion at the wing border (Bookstein, 1991).

**Fig 3.**
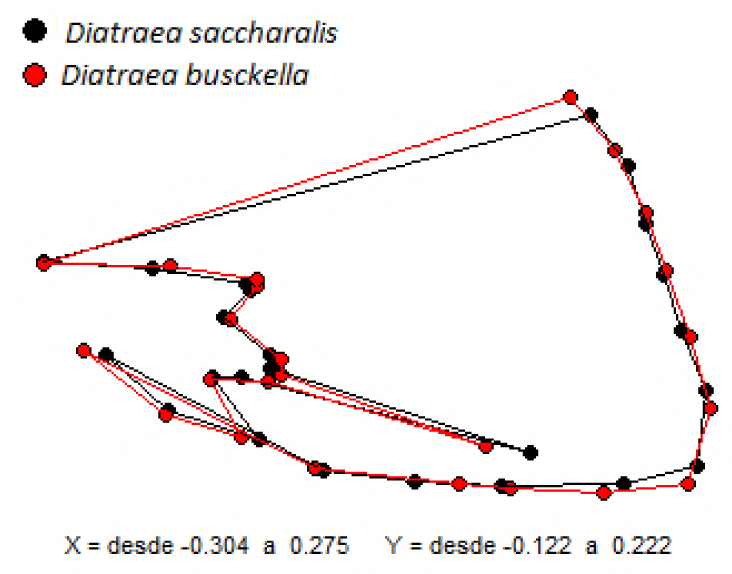
Distances and formation of 24 morphometric points for *D. saccharalis* and *D. busckella*.

**Fig 4.**
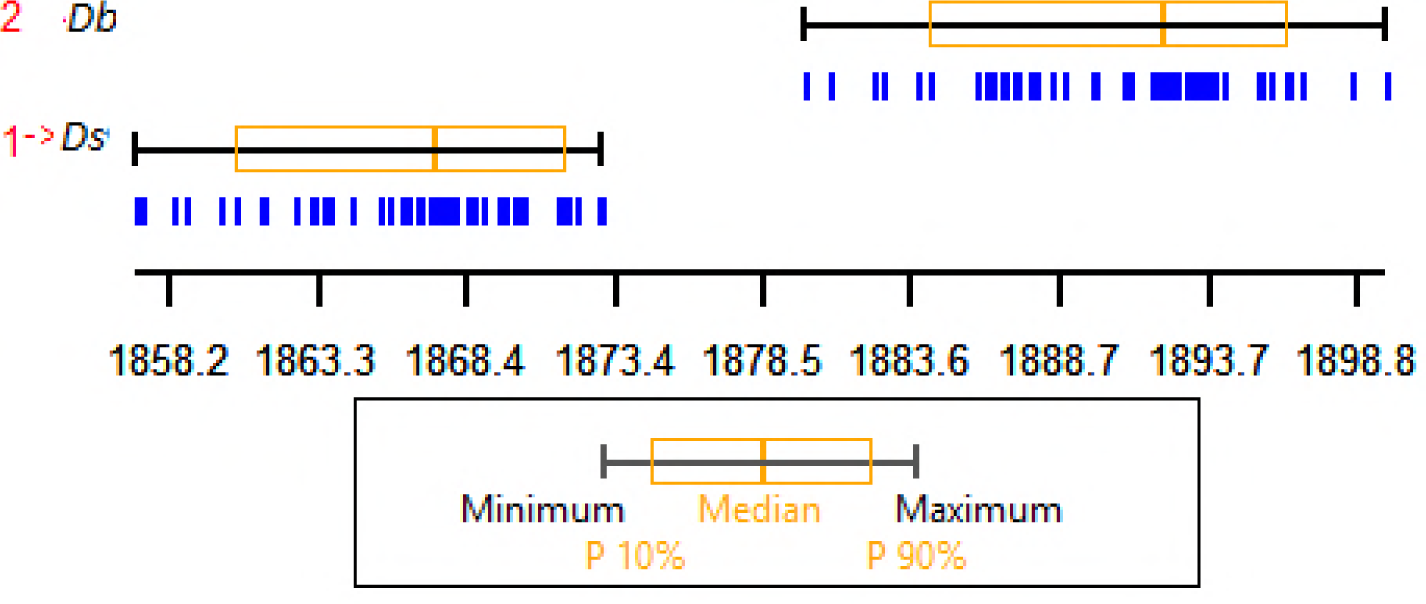
Centroid size for *Diatraea busckella* and *Diatraea saccharalis*.

Thereafter, isometric size was eliminated and morphological conformation variables were created to evaluate the relative amounts of variation between individuals, the value of asymmetry (if both sides of the specimens have been measured) and measurement error.

For the repeatability estimation (R), corresponding to two repeated measurements of the same points on 30 randomly selected images. R provides a proportion of the variance between measurements, individuals and the total variance (Henry *et al*., 2010).

The input file should contain the two successive measurements of exactly the same individuals in exactly the same order, indicating the extent to which two measurements of the same individual are correlated. To decrease the error level, 1000 permutations were run. Coordinates with high repeatability were used for the analysis. The repeatability tests were carried out using the VARwin program developed by Jean Pierre Dujardin (IRD. Montpellier, France) and freely available on the Internet at http://www.mpl.ird.fr/morphometrics/).

## Results and discussion

The 24 reference morphometric points allow to differentiate *D. saccharalis* and *D. busckella*. With distances of X from −0. 304 to 0. 275 and Y between −0. 122 to 0. 222, where both species can be separated only by generalized analysis of Procrustes (Fig 3).

### Repeatability analysis

The repeated measurements twice in order to decrease the level of error, showed an excellent relationship both in the centroid size (df= 0. 91 between species and 0. 45 between groups), and in the aligned landmarks after generalized analysis of Procrustes, being higher than 0. 9 between species (table 2). The repeatability in this case was 0. 066, indicating a large separation between the groups evaluated. In relation to the permutations, no greater distance was found in the ratio 0 / 1000 (0. 000).

**Table 2.** Generalized Procrustes analysis.

| Procrustes | Between species | Within groups |
| --- | --- | --- |
| SS | 0,04 | 0,01 |
| df* | 1,39 | 0,68 |
| MS | 0,06 | 0,01 |

At the time of the simple Re-classification, the classified individuals who were part of the discriminant analysis, presented a correct assignment, where *D. saccharalis*: 50 / 50 (100%) *D. busckella*: 50 / 50 (100%), *as second verifier*, was used the Cross Classification where *D. saccharalis*: 50 / 50 (100%) and *D. busckella*: 50 / 50 (100%), indicating that the location of the morphometric points and wings are within the correct group.

### Size

Both in the aggregate sample of 50 *D. saccharalis* and 50 *D. busckella* in each range of factors assessed, the difference between species in size was evident: *D. busckella* is significantly larger than *D. saccharalis*. The centroid size of the size variation by species was obtained as quartile graphs. Each box shows the median and quartiles 10 and 90. Below each box, the blue lines show the distribution of individuals according to their size. The units are pixels converted to millimeters, where the Mean and type of interval was 53,109; 4,071 respectively (Fig 7).

**Fig 7.**
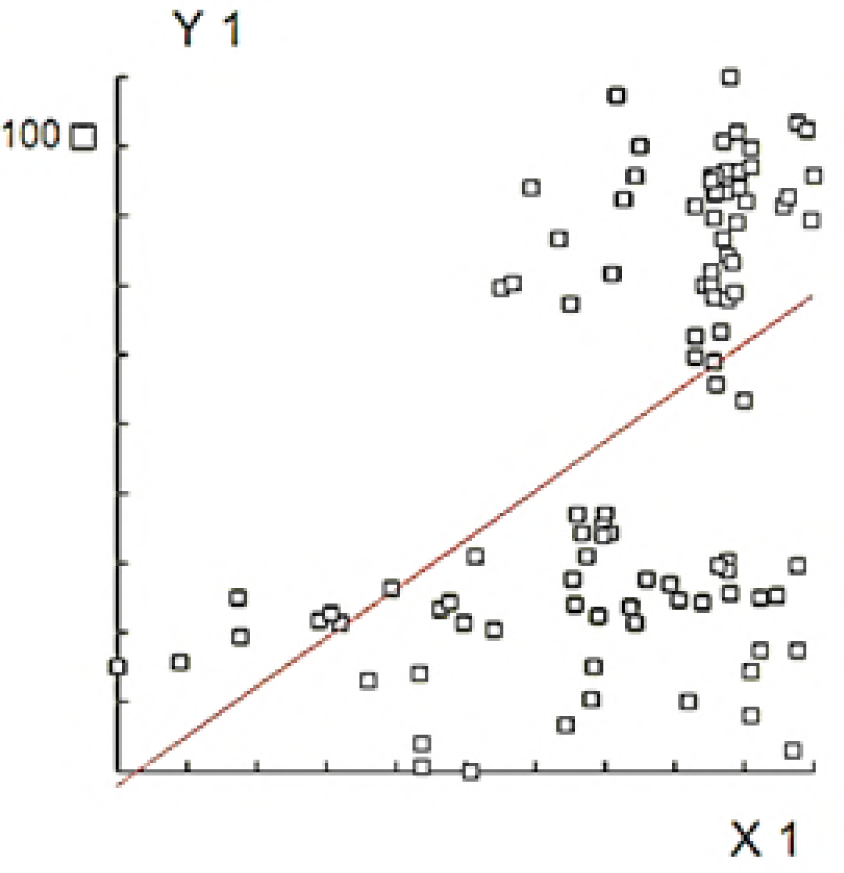
Correction of size influence on the conformation coordinates for *D. busckella* and *D. saccharalis*.

The Principal Components Analysis showed two large conglomerates according to wing shape (Fig 5). The first conglomerate was made up of *D. saccharalis* and *D. busckella*, where the contribution was PC1 0. 999997360 and PC2 0. 999998647.

**Fig 5.**
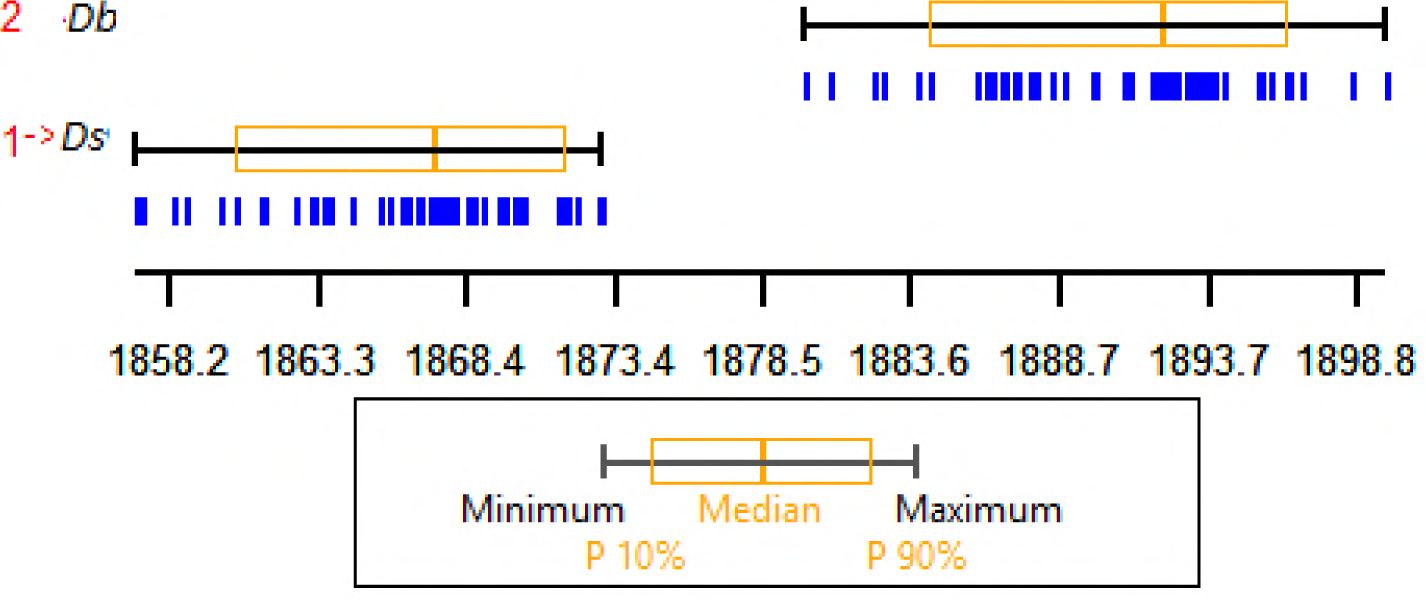
Principal component analysis. *D. busckella* and *D. saccharalis* for 50 individuals of each species.

### Alometry

In both species the alometry was highly significant (p<0. 0001). However, when the size of the individuals was removed, the wing conformation continued to be different, the alometric slopes were in the same direction (Wilks= 0. 001; df=98; r = 25%), i. e., the differences in conformation were not due to a size intervention, the conformation was free of alometry (p=0. 000) (Fig 6). However, when correcting the influence of size on the conformation coordinates, through the model of alometric slopes, the variability of shaping lingered (p<0. 0001), considering that the difference between species in conformation was free of size, these variations found can be caused by evolutionary divergence processes, more related to its acquis (r = 45%, df = 98) (Fig 7).

**Fig 6.**
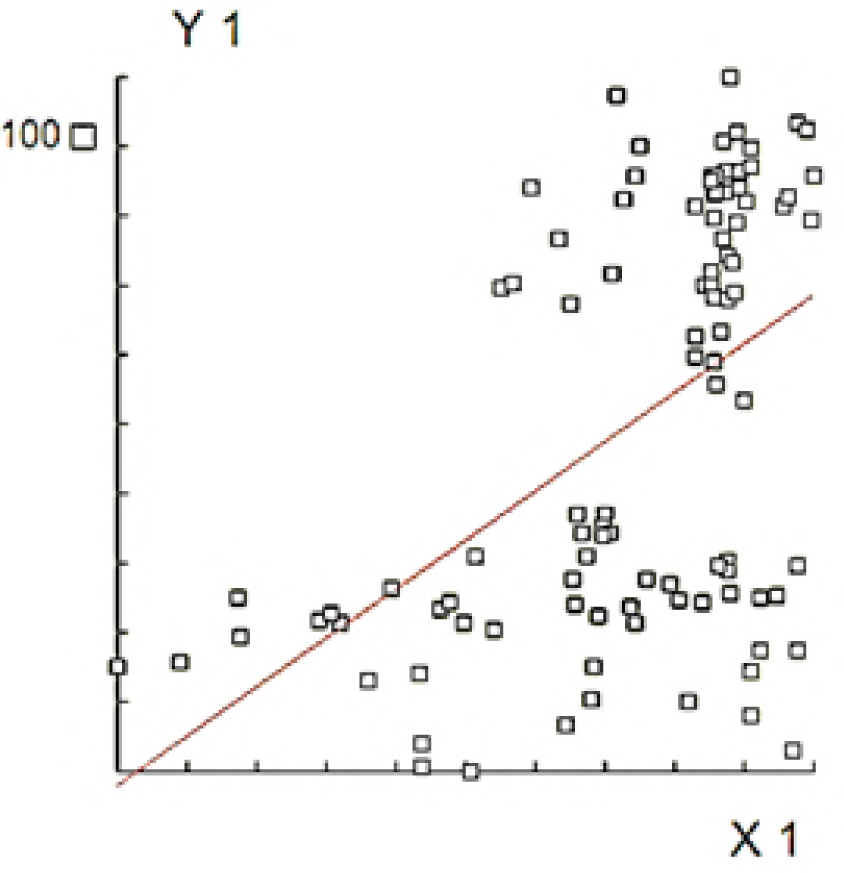
Differences in conformation without size intervention for *D. busckella* and *D. saccharalis*.

The Mahalanobis distances calculated between each pair of species were all significant (53. 56) when evaluating the null hypothesis by 1,000 permutations; showing that the 2 species collected in the different regions of Caldas are different and that these results are not random products. In addition, the centroids (multivariate conformation means) of each entity allow the species to be clearly differentiated. This demonstrates the importance of the conformation of the wing to separate the species of the genus *Diatraea*

In *S. frugiperda* Individuals differed significantly in the size and shape of their wings, and therefore in the geometric shape of their wings. Morphometry represents a promising tool for recognizing specimens of each strain of specimen collected in the field (Cañas *et al*., 2016).

### Relative wing deformations

The differences in wing shape were displayed on the deformation grids. The direction of the APRs movements was opposite between *D. saccharalis* and *D. busckella*. In both species the vectors of APRs 4-5, 12-13 and 6 were directed towards the center of the wing border, the arrows of APRs 14, 15 and 16 were directed towards the edge of the costal vein, the vector of PAR 19 upwards and the vectors of APRs 17 and 18 downwards along the border of the wing, showing a tendency to widen in the APRs located at the border of the wing (Fig 8).

**Fig 8.**
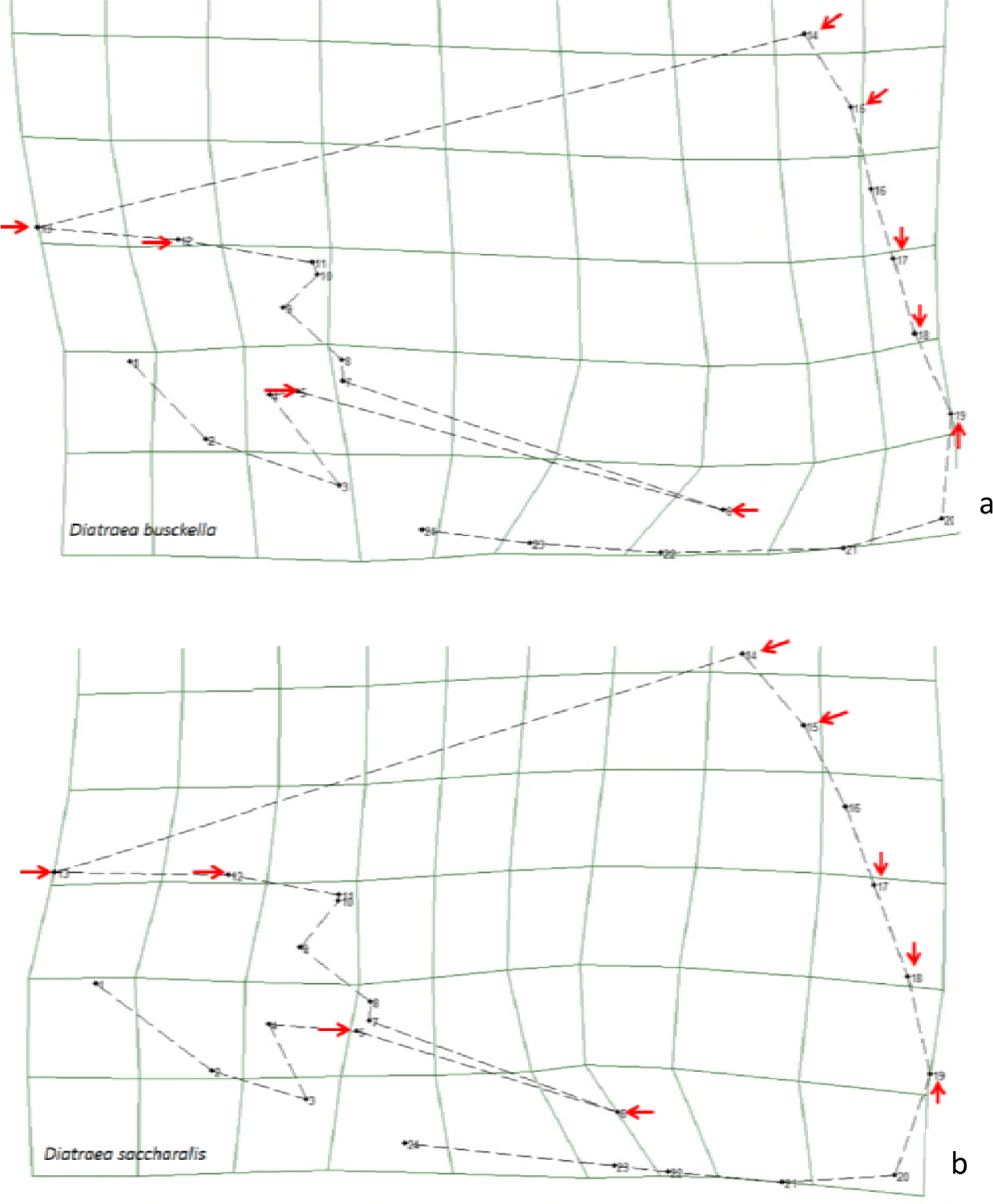
Relative wing deformation for a) *Diatraea busckella* and b) *Diatraea saccharalis*.

The contrast between *D. saccharalis* and *D. busckella* in size and conformation could be confirmed. In *Diatraea* spp the difference in size had been detected in pupae and adult states (Betts y Wootton,1988; Calle *et al*., 2008; Cadena, 2008; De Vries *et al*., 2010). However, the divergence in the wings may provide information about possible environmental and genetic pressures that species are suffering (Dellicour *et al*., 2017). Several studies show that the difference between size and habits can influence the life history traits of insects (Cadena, 2008).

#### Size

Wing size is usually associated with the size of the insect’s body (Dujardin, 2008; Francuski. *et al*., 2009). In the size of *D. saccharalis* and *D. busckella*, the difference was striking, *D. busckella* was larger than *D. saccharalis* (p<0. 0001). This trend is very common in butterflies, moths, and other insects (Adams & Funk, 1997; Hernández *et al*. 2010; Henry *et al*. 2010, Sontigun *et al*., 2017), several authors agree that they reflect different selection pressures according to the species (Ayala *et al*, 2011; Sontigun *et al*, 2017). The larger size may be favored by natural selection in the case of females to increase fecundity, suitability and lifespan (Hernández, *et al*. 2010, Francuski, *et al.,* 2009, Adams & Funk, 1997). Whereas the small size of the male may be the result of sexual selection because of the rapid development that leads to early hatching (Hernández *et al*. 2010). In *D. saccharalis* and *D. busckella* there is no work to support this statement, in females the dynamics of the species have not been measured in terms of size, but in males if early hatching has been observed with respect to females (Baena, 2010).

### Conformation

The difference of *Diatraea* spp was highly significant (p < 0. 0001) (Fig 9). Wing conformation may be affected by the two types of selection: 1) Natural, which is related to flight ability; (Francuski *et al*., 2009; Gidaszewski *et al*., 2009; Hernández *et al*., 2010), 2) sexual, related to differences in reproductive roles between males and females (Marsteller *et al*., 2009). The function in relation to flight habits and distances travelled is an important mechanism of isolation between species and wing shape can be a response to life traits and histories, this selection pressure can accelerate evolution and is associated with speciation of various groups (Andersson, 1994; Marsteller *et al.,* 2009).

**Fig 9.**
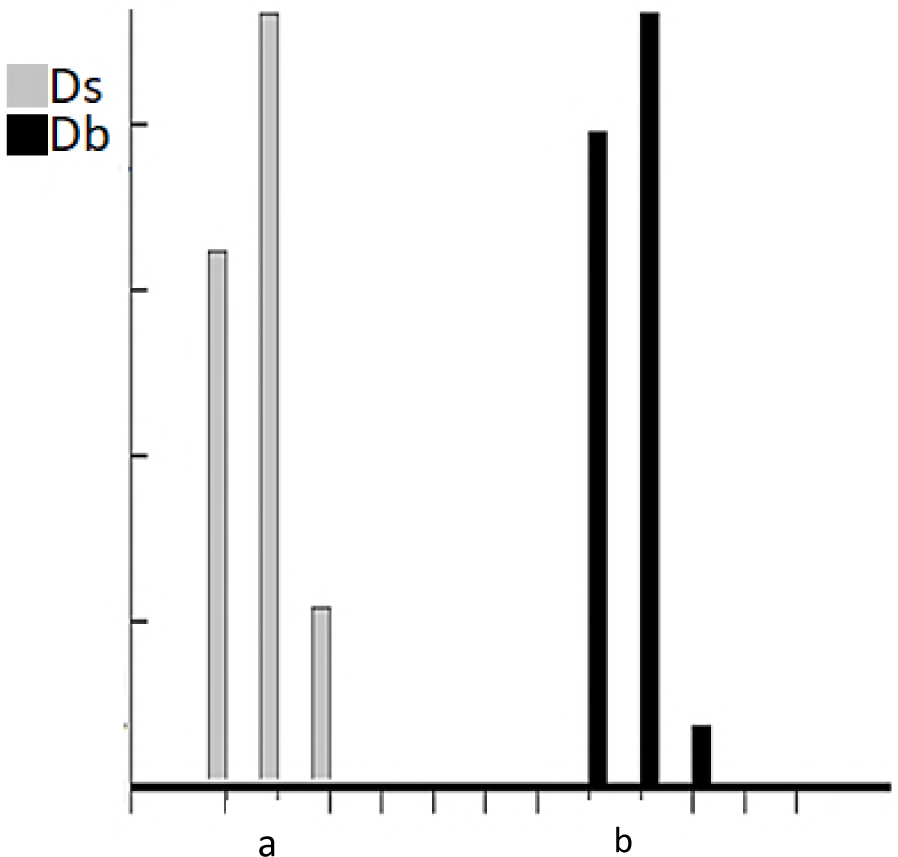
Individuals grouped by species. a). *D. saccharalis in white*, b). *D. busckella* in black) along the discriminant axis.

In *D. busckella* an elongation of the wing was noted (Figure 12), this tendency may be related to the periods of flight required to oviposit or find the female sexually active (Hernández *et al.,* 2010; Dellicour *et al*., 2017). A longer wing reduces energy requirements, breaks frictional forces and improves aerodynamic drag, giving greater dispersion ability required for long flight periods (Wotton, 1992; Betts and Wootton, 1988; DeVries *et al.,* 2010; Shi *et al.,* 2015).

In the aggregate sample, the difference in the conformation of the wings *D. busckella* and *D. saccharalis* was highly significant (Wilks = 0. 57, df = 99 p<0. 0001). As there was no overlap between the species and according to the validated Landis and Koch classification, the concordance of the reclassification was perfect (Landis & Koch, 1977; Ayala *et al.,* 2011).

## Conclusions

Morphometric tools have proven their usefulness in the discrimination of arthropods, particularly in vector insects that cannot be identified by traditional morphological characteristics, as well as in studies of intra-specfic variability (Dujardin *et al*., 1997, 1998; Rubiio-Pallis, 1998; calle *et al*., 2002, 2008; Sontigun 2017).

Geometric morphometry, describing the changes in shape from the separation of its most important components (size and configuration), open an innumerable possibilities for the performance of inter and intra-specific analyses, as well as for application in areas such as taxonomy, systematics, biomechanics, ethology and ecology (Becharia and Liria, 2012).

The shape of the wings can be a useful tool to separate the taxa at the genus and species level depending on the taxa analyzed. It can be highly reliable, especially for classifying *Diatraea* species. Alometry did not affect species separation (Sontigun *et al*., 2017).

In the present study, the importance of using geometric morphometry is noted. By including the anatomical points that defined the geometric configuration of the wing, the conformation of each species was obtained, and they were separated from each other with a high degree of confidence. In addition, it was possible to visualize the differences in wing shape between the species.

The use of geometric morphometry clearly differentiated between *D. busckella* and *D. saccharalis*, eliminating the environmental factors that could generate some level of error, being considered a support tool that validates the molecular biology processes for the identification of organisms.

It was corroborated that the presence of *D. busckella* is clear in the department of Caldas, and is in greater proportion than *D. saccharalis*, being possible the colonization of this species by the effect of global warming, it is recommended to carry out work to validate this hypothesis.

## Acknowledgements

To the National Sugar Cane Center of Colombia CENICAÑA for its valuable contribution and collaboration in the supply, development and analysis of information.

To Research Center for Research, Innovation and Technology to the Sugarcane Sector of the Caldas Department BEKDAU, for contributing significantly to the development and execution of the project.

To Washington State University WSU, for guiding me in the development and analysis of each of the parameters evaluated

